# Experimental evolution of virulence and associated traits in a *Drosophila melanogaster* – *Wolbachia* symbiosis

**DOI:** 10.1101/2020.04.26.062265

**Authors:** David Monnin, Natacha Kremer, Caroline Michaud, Manon Villa, Hélène Henri, Emmanuel Desouhant, Fabrice Vavre

**Author notes:** **Cite as:** Monnin D, Kremer N, Michaud C, Villa M, Henri H, Desouhant E, Vavre F (2020) Experimental evolution of virulence and associated traits in a *Drosophila melanogaster* – *Wolbachia* symbiosis. bioRxiv, 2020.04.26.062265, ver. 4 peer-reviewed and recommended by PCI Evol Biol. https://doi.org/10.1101/2020.04.26.062265u.

## Abstract

Evolutionary theory predicts that vertically transmitted symbionts are selected for low virulence, as their fitness is directly correlated to that of their host. In contrast with this prediction, the *Wolbachia* strain *w*MelPop drastically reduces its *Drosophila melanogaster* host lifespan at high rearing temperatures. It is generally assumed that this feature is maintained because the *D. melanogaster–w*MelPop symbiosis is usually not exposed to environmental conditions in which the symbiont is virulent. To test this hypothesis, we submitted *w*MelPop-infected *D. melanogaster* lines to 17 generations of experimental evolution at a high temperature, while enforcing late reproduction by artificial selection. The fly survival was measured at different time points, as well as two traits that have been proposed to be causally responsible for *w*MelPop virulence: its relative density and the mean number of copies of octomom, an 8-genes region of the *Wolbachia* genome. We hypothesised that these conditions (high temperature and late reproduction) would select for a reduced *w*MelPop virulence, a reduced *w*MelPop density, and a reduced octomom copy number. Our results indicate that density, octomom copy number and virulence are correlated to each other. However, contrary to our expectations, we could not detect any reduction in virulence during the course of evolution. We discuss the significance of our results with respect to the evolutionary causes of *w*MelPop virulence.

## Introduction

Symbionts live inside (endosymbionts) or on (ectosymbionts) bigger organisms referred to as hosts. As a result, their evolutionary success crucially depends on their ability to colonize, or be transmitted to, new hosts. Deleterious symbionts, also known as parasites, exploit their hosts in a way that maximizes their transmission, but usually face a trade-off between instantaneous transmission rate and opportunities for transmission. Indeed, the reproductive rate of a parasite is likely to be positively correlated to both its instantaneous transmission rate and its virulence (the reduction in fitness it incurs to its host). Increased virulence is in turn associated, notably through increased host mortality, with a reduction in the opportunities for transmission (Anderson & May 1982, Ewald 1983, see Alizon et al. 2009 for a review and defence of this hypothesis).

Theoretical and experimental works suggest that the mode of transmission – which ranges from fully vertical (from parents to offspring) to fully horizontal (contagion) – is an important factor shaping parasite virulence. The fitness of vertically transmitted symbionts being tightly correlated to that of their hosts, they are expected to be selected for low virulence (Ewald 1983). This reasoning, just like the transmission/virulence trade-off hypothesis, rests solely on the inter-host level of selection (*i.e.,* selection between symbionts infecting different hosts), ignoring the potential effects of intra-host selection (*i.e.*, selection between symbionts infecting the same host). Indeed, the most competitive symbionts within a host may replicate faster and be more virulent than what would be optimal from the standpoint of inter-host selection (Alizon et al. 2013). However, the causal link from vertical transmission to low virulence is well supported by the experimental evolution of parasites following the enforcement of different transmission modes (Bull et al. 1991, Turner et al. 1998, Messenger et al. 1999, Stewart et al. 2005). Accordingly, the most widespread vertically-transmitted symbiont in insects, the intracellular bacterium *Wolbachia,* is often found to be either avirulent (Hoffmann et al. 1994, Giordano et al. 1995, Bourtzis et al. 1996, Poinsot & Merçot 1997, Hoffmann et al. 1998) or slightly virulent (Hoffmann et al. 1990, Turelli & Hoffmann 1995, Clancy & Hoffmann 1997).

The most striking exception to this pattern is *w*MelPop, a *Wolbachia* strain hosted by *Drosophila melanogaster* that drastically reduces its host lifespan at high rearing temperatures (Min & Benzer 1997). This exceptional virulence raises the question of its causes, both proximal and ultimate. Min & Benzer (1997) suggested that the proximal cause of the virulence of *w*MelPop is its over-replication, which was further supported by subsequent studies (McGraw et al. 2002, Strunov et al. 2013). Furthermore, the repetition of an eight-gene region (“octomom”), has recently been proposed as the genomic basis of *w*MelPop high density and virulence (Chrostek & Teixeira 2015). Octomom includes genes encoding proteins potentially involved in DNA replication, repair, recombination, transposition or transcription (Chrostek et al. 2003) and was found to be correlated with *w*MelPop density and virulence (Chrostek & Teixeira 2015). However, both the link between octomom copy number and density and the link between density and virulence have been called into question. Rohrscheib et al. (2016) indeed found that differences in survival between flies reared at 24°C and 29°C cannot be explained by differences in bacterial density. Similarly, comparing flies of the same age but reared at different temperatures, or of different ages but reared at the same temperature, they found no correlation between octomom copy number and either density or virulence. However, to exclude the possibility that octomom copy number has an effect on density, and density on virulence, the effect of these variables should be assessed independently of temperature and age. The ultimate cause of *w*MelPop virulence has been assumed to be non-adaptive: as this *Wolbachia* strain is not known to occur in nature but may have originated in the lab, and as flies are typically reared at temperature lower than those at which *w*MelPop is virulent and reproduce early in the lab, there is no selective pressure for reduced virulence (Reynolds et al. 2003). It was indeed found that *w*MelPop reduces fly survival at 25°C, but not at 19°C (Reynolds et al. 2003). If this explanation is correct, raising *w*MelPop-infected flies at a high temperature, while enforcing late reproduction, should select for: (i) a reduced *w*MelPop virulence, (ii) a reduced *w*MelPop density (under the assumption that *w*MelPop virulence at high temperature is due to its high density), and (iii) a reduced number of octomom copies (under the assumption that *w*MelPop virulence at high temperature is due to its high octomom copy number).

In the present study, we used experimental evolution in conjunction with artificial selection over 17 generations to test these hypotheses, raising *w*MelPop infected flies at 29°C, and enforcing reproduction eight days after emergence. In addition, in one experimental condition, we fed the flies with paraquat (1,1’dimethyl-4-4’bipyridynium dichloride), a pro-oxidant compound and herbicide that was previously shown to increase the survival of *w*MelPop-infected flies (while having no effect on the survival of uninfected flies) and to reduce the symbiotic density (Monnin et al. 2016). We hypothesised that this treatment would reduce or cancel the selective pressure for reduced virulence, density, and octomom copy number, resulting in higher values for these parameters, compared to the control condition. The results of these experiments were interpreted in light of our assessment of the relationship between octomom copy number and the age of the host. Despite not being known to occur in nature, the *D. melanogaster-w*MelPop association can help to understand the real-life dynamics of *host-Wolbachia* associations, especially in their early stages. Indeed, one can expect that the low virulence typically observed in *Wolbachia* symbioses is the outcome of a relatively long process of coevolution between the symbiotic partners. Newly formed associations, by contrast, might be more likely to exhibit levels of virulence similar to what is observed in *D. melanogaster-w*MelPop associations. Furthermore, our results contribute to the ongoing debate concerning the relationships between *w*MelPop octomom copy number, density, and virulence (Rohrscheib et al. 2016, Chrostek & Teixeira 2017, Rohrscheib et al. 2017).

## Methods

### Biological material

We used 17 isofemale lines of *w*MelPop-infected *D. melanogaster*^w1118^. These lines, originated from flies provided by Scott O’Neill (Monash University, Australia), were founded approximately six months before the start of the experiment, and then maintained under controlled rearing conditions at 18°C (12 LD cycle), a temperature at which *w*MelPop is expected to be avirulent (Reynolds et al. (2003) found no effect of the infection on mortality rate at 19°C and no virulence of *w*MelPop was ever detected at a lower temperature). The flies were maintained on standard drosophila medium (David & Clavel 1965).

### Experimental evolution procedures

The experiment was conducted at 29°C (12 LD cycle), a temperature at which *w*MelPop is known to be virulent (Min & Benzer 1997, Strunov et al. 2013, Monnin et al. 2016). Emerging flies from the 17 isofemale lines were mixed to constitute the founding generation G0 and were allowed to lay eggs. Twenty experimental lines were then constituted, each originating from 100 randomly-picked eggs that were deposited in vials containing 1.5 g of *Drosophila* medium. The experimental lines thus constituted were split into 10 “paraquat lines”, in which larvae developed on drosophila medium supplemented with 150 μL of a 10 mM paraquat (1,1’dimethyl-4-4’bipyridynium dichloride) solution (as described in Monnin et al. 2016), and 10 “water lines”, in which the paraquat was replaced by 150 μL of water. The G1 flies emerging from each vial were then allowed to lay their eggs on unsupplemented medium for eight days (the medium was not supplemented with paraquat or water, as this would have been a confounding factor for the subsequent measurements). The medium was changed regularly, including at the end of day seven, and the eggs already laid on it were therefore removed. For each experimental line, a subset (n=100) of the eggs laid on the eighth day following emergence was deposited in new vial similar to those in which the parents developed – *i.e.,* paraquat-supplemented medium for the paraquat lines and water-supplemented medium for the water lines. The protocol was repeated for the emerging G2 flies, and again until G17. The evolution of *w*MelPop virulence, density and mean octomom copy number was measured as follow: in each line, eggs laid seven days after emergence were collected at G2, G3, G9, G12 and G17 and allowed to develop on unsupplemented medium. Then, emerging females were used either to assess their survival, or their *Wolbachia* density and mean octomom copy number (fig. 1). These traits were not measured at every single generation, as we do not expect the changes from one generation to the next to be so great as to justify measuring all traits at each generation.

**Figure 1.**
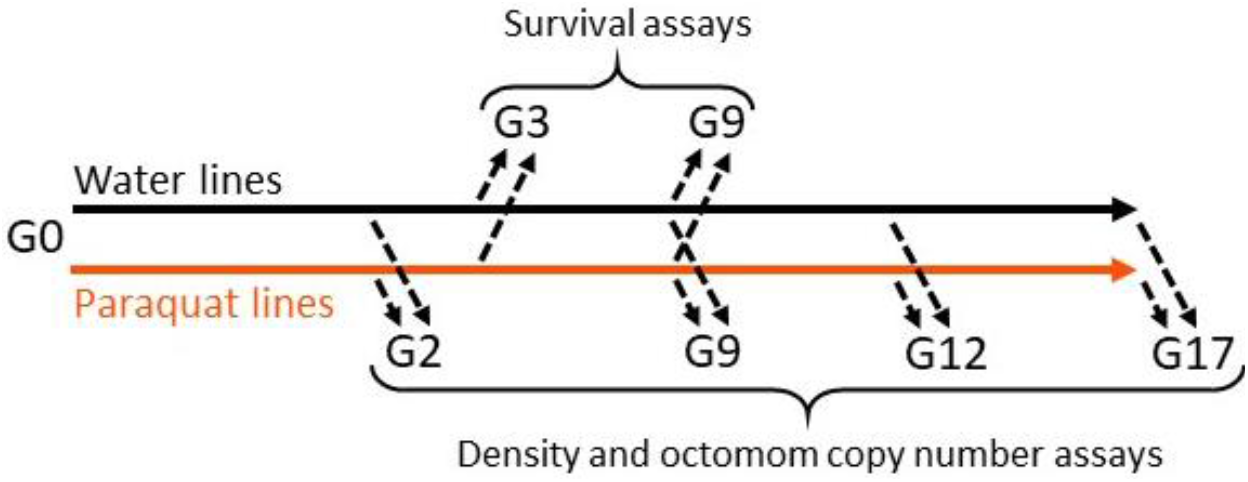
Summary diagram of the experimental evolution experiment, showing the generations (G0-G17) at which the different measurements were performed.

Though we chose – based on preliminary observations – the eight days period to enforce late reproduction while reducing the risk of line extinction, several lines nevertheless went extinct before the end of the experiment, as a result of lack of egg laying. Three control lines and two paraquat lines went extinct between G3 and G9, four control lines and one paraquat line between G9 and G12, and one control line and two paraquat lines between G12 and G17.

### Survival assays

Fly survival was measured, as a proxy for *w*MelPop virulence, at G3 and G9. At emergence, 15 females from each experimental line were dispatched into three vials containing sugar-supplemented agar (10% sugar). The surviving flies were then counted every day. The line extinction rate – the ratio of lines that went extinct over the total number of lines – was calculated for both the water and the paraquat lines, both during the 8 generations prior to G9 and during the 9 subsequent generations. It serves as another proxy for *w*MelPop virulence. We used Fisher’s exact tests to assess differences in extinction rates.

### DNA extraction

The *w*MelPop density and mean octomom copy number were measured at G2, G9, G12 and G17. For each generation and line, four of the emerging females were transferred on sugar-supplemented agar (10% sugar), to be collected four days later and then kept at −80°C. Because of technical issues during DNA extraction or qPCR, the final number of replicates varies from 2 to 4 (mean number > 3.8). The DNA extractions were performed using the 96 wells Biobasic EZ-10 BS4372 kit. The entire flies were mechanically crushed for 30 s at 25 Hz using a 5-mm stainless steel bead in a TissueLyser (Qiagen). Three hundred μL of ACL solution and 20 μL of 16 g.L^-1^ proteinase K were then added to the samples. Following an overnight incubation at 56°C, 300 μL of AB solution was added. The samples were then transferred on the plate from the kit, and the purification was performed following the instructions of the manufacturer. The DNA was eluted in 100 μL of elution buffer and stored at −20°C.

### Quantitative PCR

Both relative *w*MelPop density and mean octomom copy number were measured in each DNA extract by quantitative PCR, conforming to the Minimum Information for Publication of Quantitative real-time PCR Experiments (MIQE) guidelines (Bustin et al. 2009). The relative *w*MelPop density was quantified by amplifying two monocopy genes: one *Wolbachia* gene (*wd0505)* and one *D. melanogaster* gene (*rp49).* The mean octomom copy number was quantified by amplifying two *Wolbachia* genes: one that is part of the octomom region (*wd0513),* and one that is not (*wd0505*). The primers used to amplify *rp49* were as follow: *rp49dd-F:* 5’-CTG-CCC-ACC-GGA-TTC-AAG-3’, *rp49dd-R:* 5’-CGA-TCT-CGC-CGC-AGT-AAA-C-3’. The primers used to amplify the *wd0505* and *wd0513* genes were published in Chrostek et al. (2013). The primers were synthetized by Eurogentec (Seraing, Belgium). qPCR mixes consisted of 5 μL of Sso Advanced SYBR Green Supermix (Bio-Rad, Hercules, California, USA), 1 μL of water, 0.5 μL of each primer (10 mM), and 3 μL of DNA samples diluted at 1/10. The reaction conditions for amplification were 95°C for 3 min, followed by 40 cycles of 95°C for 10 s, 59°C for 10 s and 68°C for 15 s. A melting curve was recorded from 70°C to 95°C to ensure the specific amplification of the transcript. Amplification and detection of DNA by quantitative PCR were performed with CFX96 instrument (Bio-Rad). Standard curves were plotted using seven dilutions (10^2^ to 10^7^ copies) of a previously amplified PCR product that had been purified using the kit Nucleospin extract II (Machery-Nagel). Primer sets exhibited mean efficiencies of 93.2% ± 1.2 for *rp49dd,* 91.0% ± 0.7 for *wd0505,* and 92.7% ± 0.6 for *wd0513.* Two technical replicates per biological replicate were used for the determination of DNA starting quantities. As the deviations between these duplicates were below 0.5 cycles, the mean Cp values were calculated. As PCR efficiency were close to 100%, we calculated the relative *w*MelPop density and the mean octomom copy number using the following formulas: *2^Cp_rp49-Cp-wd0505^* and *2^Cp_wd0505-Cp_wd0513^*, respectively.

### Relationship between mean octomom copy number and the age of the host

To evaluate the possibility that intra-host selection is involved in the evolution of *w*MelPop octomom copy number, we assessed the relationship between the mean octomom copy number and the age of the host. To do so, we performed qPCR as described above on samples obtained in the course of a previously published experiment (Monnin et al. 2016). Briefly, *w*MelPop-infected *D. melanogaster*^*w*1118^ were raised until emergence on standard drosophila medium supplemented with water or paraquat (as described above). Emerging females were either immediately collected and kept at −80°C (1 fly per treatment) or transferred on sugar-supplemented agar (10% sugar) to be collected at days 7, 14 and 21 of adulthood (four flies per day and per treatment). DNA extractions were performed as described above.

### Statistical analyses

We used the R software (version 3.3.2) for all analyses (R Core Team 2016).

We tested for evolution in survival between G3 and G9 using a mixed generalized linear model with a gamma distribution (in the absence of censored data, a survival model is not necessary, and the gamma distribution is suitable for continuous positive data). Individual longevity was used as the response variable. Generation (G3 or G9) and treatment (water or paraquat) were treated in the model as fixed factors, while the line and vial factors were included as random effects.

The evolution of density and mean octomom copy number was tested using mixed linear models, with the generation included as a fixed quantitative variable, the treatment as a fixed qualitative variable, and the line factor included as a random effect. Normality and homoscedasticity were checked graphically for both models.

The relationship between *Wolbachia* density and survival at G9 was tested by fitting a linear model with median longevity as the response variable, mean density as a quantitative explanatory variable, and the treatment (water or paraquat) as an explanatory factor.

The relationship between density and mean octomom copy number was tested by fitting a mixed linear model with density as the response variable, mean octomom copy number as a fixed quantitative explanatory variable, and generation, line and treatment as random effects.

The relationship between age and mean octomom copy number was tested by fitting a linear model with mean octomom copy number as the response variable, age as a quantitative explanatory variable and treatment as a qualitative explanatory variable.

## Results & Discussion

### Test of the predictions

We followed the evolution of *w*MelPop virulence, density, and octomom copy number at a high temperature while enforcing late reproduction. Our first prediction was that during the experimental evolution, *w*MelPop virulence would have decreased in the water lines, and less so, or not at all, in the paraquat lines. To test this, we compared fly survival at G3 and G9. We can be confident that fly mortality is induced by *w*MelPop, as the strong reduction of host longevity induced by *w*MelPop has been established previously. A study performed in our lab (Monnin et al. 2016), using the same *D. melanogaster* line and the same survival protocol, found a large mean difference in longevity (4.4±0.66 days) between *w*MelPop-infected and uninfected individuals.

Survival data at G3 confirmed that mortality began before the eighth day of the adult life of the flies (fig. 2a), suggesting that the late reproduction we enforced should have induced a selective pressure for decreased *w*MelPop virulence, at least in the water lines. In the paraquat lines, we expected a lesser reduction, or no reduction at all, in virulence.

**Figure 2.**
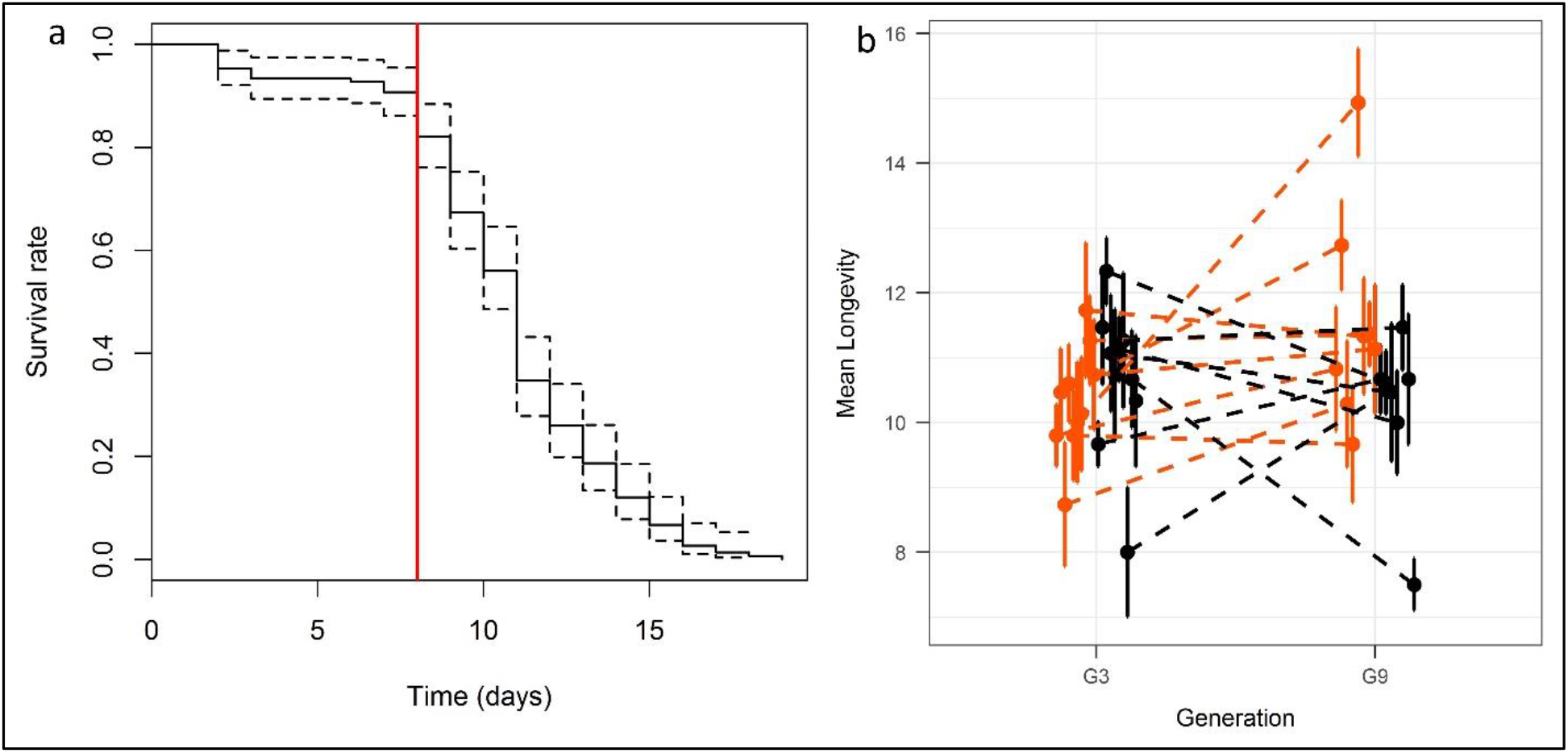
Evolution of fly survival. (a) Survival of female flies in the water condition at G3. Dotted lines represent the 95 % confidence interval. The red vertical line shows the age (8 days) at which reproduction was enforced during the experimental evolution. The corresponding figure for the paraquat condition is provided as supplementary material (b) Mean longevity (± standard error) in all lines in the paraquat (orange) and water (black) conditions, at generations 3 and 9.

Neither the interaction between generation and treatment (χ^2^_1_=3.23, p=0.07) nor the treatment effect (χ^2^_1_=0.38, p=0.54) was statistically significant, suggesting that the paraquat treatment did not modify the evolution of survival, as compared to the water condition. Furthermore, there was no effect of the generation factor, indicating that survival did not increase between G3 and G9 (χ^2^_1_=0.98, p=0.32; fig. 2b; median longevity at G3 in the control lines: 11 days; at G9 in the control lines: 10.5 days; at G3 in the paraquat lines: 10 days; at G9 in the paraquat lines: 11 days). As the survival was last assessed at G9, we cannot directly test whether it evolved between G9 and G17. Given that an increased survival would presumably lead to a decreased probability of extinction, we compared the rates of line extinction per generation before and after G9. In the water condition, it was 0.038 before G9 (3 out of 10 lines went extinct at some point during these eight generations), and 0.079 after G9 (5 out of 7 lines went extinct at some point during these nine generations), showing that no increase in extinction rate occurred during the experimental evolution, even after G9. Likewise, in the paraquat condition, the rate of line extinction per generation was 0.025 before G9 (2 out of 10 lines went extinct at some point during these eight generations), and 0.042 after G9 (3 out of 8 lines went extinct at some point during these nine generations). Differences in extinction rate between the early and late generations were not significant (Fisher’s exact tests, p=0.15 in the water lines, p=0.61 in the paraquat lines, p=0.16 overall).

Taken together, these survival and extinction rates suggest the contrary to our prediction, the virulence of *w*MelPop did not decrease during the experiment and was not differentially affected by the treatment (paraquat or water).

The discrepancy between the first prediction and the results invalidates the next two predictions, as the expected reduction in *Wolbachia* density was supposed to causally explain the expected reduction in virulence, and the expected reduction in mean octomom copy number was supposed to causally explain the expected reduction in density. Accordingly, we observed neither a decrease in density nor in mean octomom copy number during the experimental evolution. On the contrary, both density (χ^2^_1_=27.84, p<0.001; fig. 3a) and mean octomom copy number (χ^2^_1_=16.95, p<0.001; fig. 3b) increased (mean density in the water lines at G2: 11.73; G9: 26.33; G12: 21.05; G17: 20.50; in the paraquat lines at G2: 13.05; G9: 12.74; G12: 18.59; G17: 33.23; mean octomom copy number in the water lines at G2: 2.50; G9: 3.45; G12: 2.66; G17: 3.43; in the paraquat lines at G2: 2.84; G9: 2.67; G12: 4.27; G17: 3.97). Neither the interactions between generation and treatment (density: χ^2^_1_=0.52, p=0.47; octomom copy number: χ^2^_1_=1.21, p=0.27) nor the treatment effects (density: χ^2^_1_=0.44, p=0.51; octomom copy number: χ^2^_1_=0.44, p=0.51) were statistically significant.

**Figure 3.**
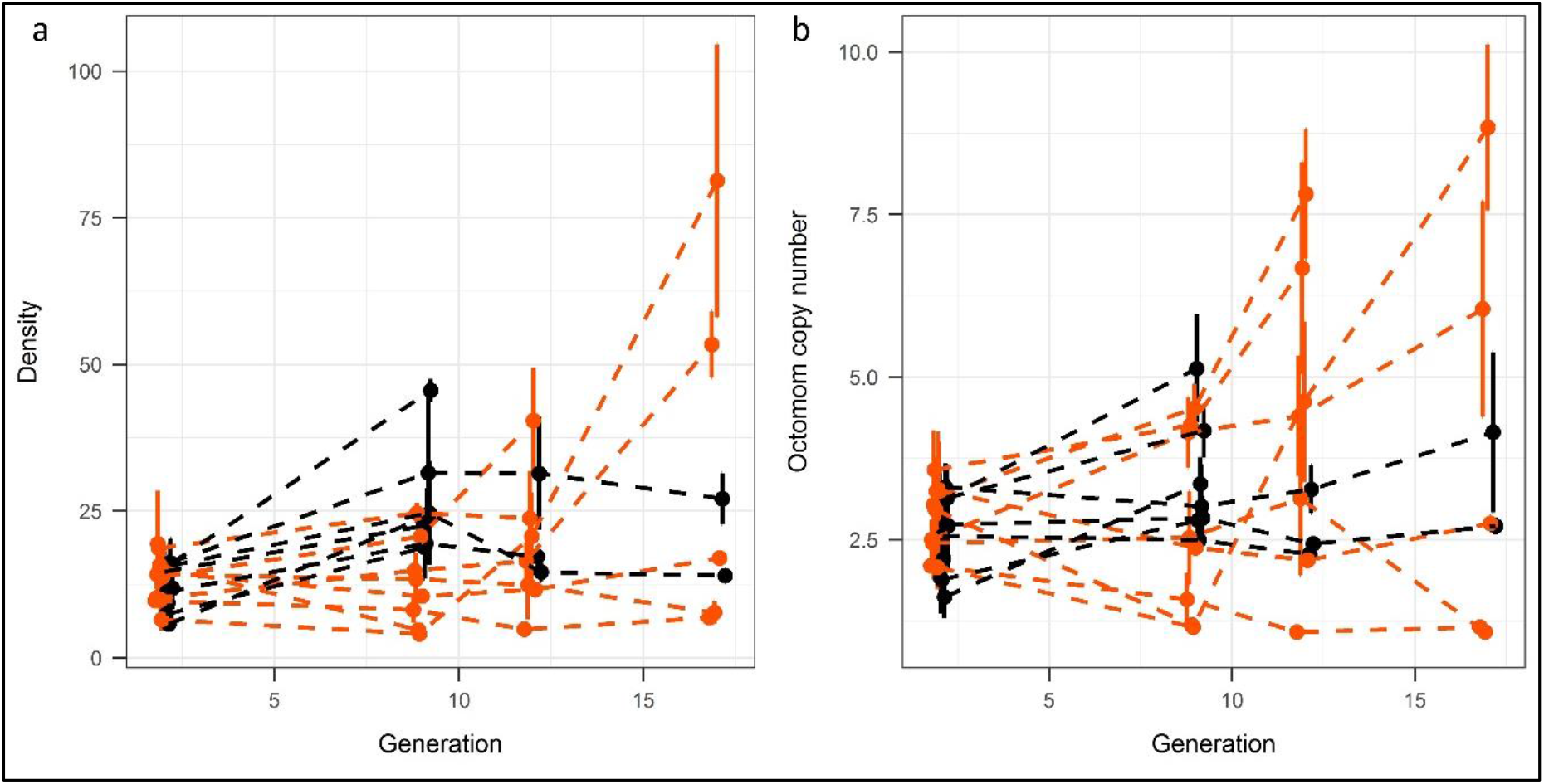
Evolution of *Wolbachia* density and octomom copy number. (a) Mean *Wolbachia* relative density (± standard error) in all lines in the paraquat (orange) and water (black) conditions, at generations 2, 9, 12 and 17. (b) Mean octomom copy number (± standard error) in all lines in the paraquat (orange) and control (black) conditions, at generations 2, 9, 12 and 17.

### Potential explanations for the falsification of the predictions

The falsification of our predictions is especially puzzling, as our results show that both density and octomom copy number evolved, suggesting that it is not by lack of time or heritable variability that the *D. melanogaster*–*w*MelPop interaction did not evolve in the expected ways.

As some lines went extinct before the end of the experiment, our statistical analyses may have been biased if those extinctions were not random. However, it is unlikely to explain why our predictions were not vindicated. Indeed, although a reduction in virulence could have been concealed by the preferential extinction of lines exhibiting low virulence, this pattern is the opposite of what we would expect. Furthermore, the observed increase in relative *Wolbachia* density and mean octomom copy number is not likely to be due to a mere increase in genetic similarity: at G2, no measured fly had a relative density superior to 40, contrary to 21% of the G17 flies (the maximum being 143); similarly, no flies at G2 had a mean octomom copy number over 6, contrary to 18% of the flies measured at G17 (the maximum being 12). Finally, we did not vary the experimental temperature – a factor recently shown to impact on the evolution of *D. melanogaster-Wolbachia* associations by Mazzucco et al. (2020) – and used only one host genetic background (*D. melanogaster*^w1118^) and one *Wolbachia* strain (*w*MelPop). We cannot exclude the possibility that different conditions would have led to different coevolutionary outcomes.

Among our starting assumptions was that the virulence of *w*MelPop is due to its high density, which in turn is due to its high octomom copy number. It is therefore noteworthy that both assumptions were recently called into question (Rohrscheib et al. 2016; see also Chroskek & Teixeira 2017, Rohrscheib et al. 2017). If, as claimed by Rohrscheib et al. (2016), the virulence of *w*MelPop is independent of its density and octomom copy number, our second and third predictions do not hold. Although this would not explain why virulence did not decrease during the experimental evolution, it would remove our puzzlement regarding the increase in *Wolbachia* density and octomom copy number. However, our results do not support Rohrscheib et al. (2016) findings. First, *w*MelPop density and fly survival were negatively related at G9, the only generation at which both survival and density data were collected (F_1,18_=5.25, p=0.04; fig. 4a). Second, *w*MelPop density and mean octomom copy number were found to be positively related (χ^2^_1_=116.31, p<0.001; fig. 4b).

**Figure 4.**
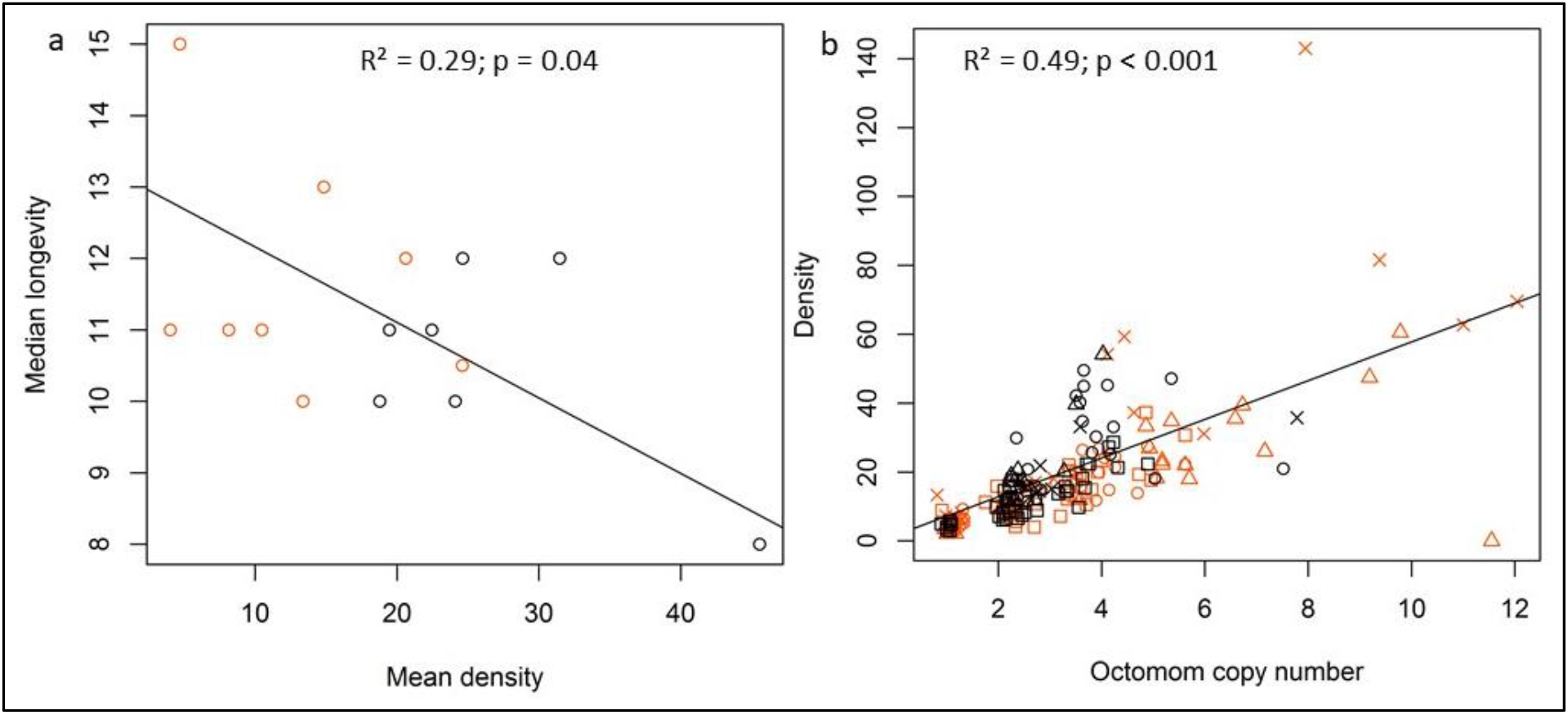
(a) Correlation between *Wolbachia* density and fly survival. Each circle represents a line in the paraquat (orange) or water (black) condition, at G9. The black line represents the regression of median longevity on mean density (data for both conditions are pooled, as the treatment effect was found not to be statistically significant). Statistics from Pearson’s correlation test are indicated. (b) Correlation between *Wolbachia* density and octomom copy number. Each symbol represents an individual fly, from the paraquat (orange) or water (black) condition, at G2 (squares), G9 (circles), G12 (triangles), or G17 (crosses). The black line represents the regression of density on octomom copy number. Statistics from Pearson’s correlation test, with all conditions pooled together, are indicated.

It is therefore more likely that our assessment of the evolutionary forces at play, rather than our assumptions regarding the proximal causes of *w*MelPop virulence, was mistaken. The increase, over time, in both density and octomom copy number, appears to be consistent with two non-mutually exclusive hypotheses. First, this pattern could be explained by the relaxation of an unknown selective pressure maintaining a low octomom copy number in flies kept at relatively low temperatures without enforcement of late reproduction, perhaps amplified by a mutational bias favouring a high octomom copy number. As each new generation of experimental evolution was started with only 100 eggs, genetic drift could have been increased by the experimental evolution/selection protocol, possibly limiting the efficiency of natural selection. However, it remains difficult to see why the selective pressure in favour of a low octomom copy number should be weaker in the experimental evolution setting than in the maintenance conditions, as our current knowledge of *w*MelPop suggests that the opposite is true: in the absence of *w*MelPop-induced mortality in the maintenance conditions, the octomom copy number is not expected to be under selection.

A second explanation for the observed pattern is that it results from directional selection in favour of high density, high octomom copy number variants, more or less constrained, in each line, by the amount of available heritable variation. Other things being equal, flies harbouring low density, low octomom copy number should be longer-lived and therefore have more opportunities to transmit their symbionts to their offspring than their counterparts infected by high density, high octomom copy number variants. As a result, inter-host selection is expected to favour *w*Melpop variants that do not reach densities so high that they compromise the fitness of their host. However, intra-host selection is also likely to affect the evolution of virulence (Alizon et al. 2013). Indeed, one can expect high density, high octomom copy number variants to replicate more rapidly than (and therefore outnumber) their low density, low octomom copy number counterparts, which will make them more likely to be transmitted to their host offspring. Crucially, the competitive advantage of rapidly-replicating variants will increase with host age, as it is positively correlated with the number of *w*MelPop generations. (At 29°C, the doubling time of *w*MelPop was estimated by Duarte et al. (preprint) to vary from 0.88 to 1.38 days.) ‘Late reproduction’ may therefore allow intra-host selection to be more powerful than inter-host selection. This is especially likely given that survival was still high at eight days (fig. 2a), suggesting that inter-host selection may have been relatively weak. Chrostek & Teixeira (2018) provide evidence for such intra-host selection by showing that the mean octomom copy number increases as flies age and reach a plateau by day eight of adulthood. Our own results confirm the increase (F1,24=25.64, p<0.001), but no plateau was observed (fig. 5). These results show that intra-host variability can exist, and therefore allow for intra-host selection, despite the bottleneck experienced by vertically transmitted symbionts (estimated to range from 850 to 8000 cells in aphid-*Buchnera* symbioses (Mira & Moran 2002)).

**Figure 5.**
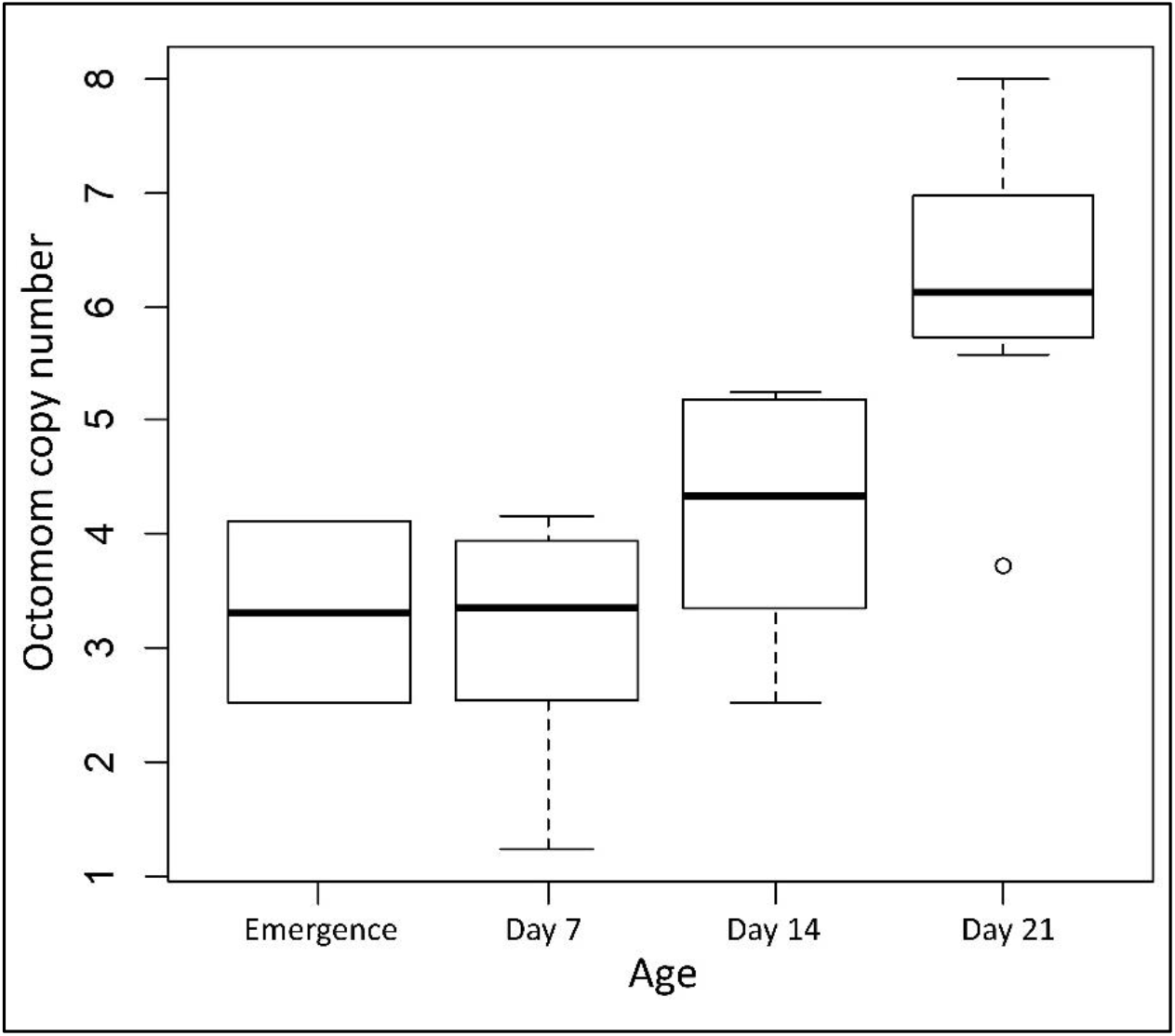
Relationship between the age of the fly and the octomom copy number of *w*MelPop. Data pooled from two treatments (paraquat and water), as neither the interaction between age and treatment (F_1,22_=2.79, p=0.11) nor the treatment effect (F_1,23_=0.14, p=0.72) were significant.

More research is needed to determine the evolutionary consequences of this intra-host dynamic. Rapid evolution of octomom copy number in *w*MelPop suggests that intra-host selection could play an important role in the evolution of vertically transmitted symbiont and may contribute to explain the persistence of relatively virulent *Wolbachia* strains in nature.

Taken together, our results show that *Wolbachia* traits associated with virulence can evolve rapidly following a switch in selection regime. These evolutionary changes challenge common assumptions on the evolution of virulence in vertically transmitted symbionts, as traits positively correlated with virulence were found to evolve. This could indicate that the intra-host level of selection plays a significant, yet underexplored, role in shaping associations with vertically transmitted symbionts.

## Data accessibility & Supplementary material

Raw data, scripts and additional figure are available online: https://doi.org/10.5281/zenodo.4065517

## Acknowledgements

We thank Nicole Lara for providing the *Drosophila* media. This work was supported by the Agence Nationale de la Recherche: ‘ImmunSymbArt’ ANR-2010-BLAN-170101 and ‘RESIST’ ANR-16-CE02-0013-01. This work was performed within the framework of the LABEX ECOFECT (ANR-11-LABX-0048) of Université de Lyon, within the programme ‘Investissements d’Avenir’ (ANR-11-IDEX-0007).

Version 4 of this preprint has been peer-reviewed and recommended by Peer Community In Evolutionary Biology (https://doi.org/10.24072/pci.evolbiol.100111).

## Conflict of interest disclosure

The authors of this preprint declare that they have no financial conflict of interest with the content of this article. NK & FV are recommenders for PCI EvolBiol.

## Notes

https://doi.org/10.5281/zenodo.4065517

### Competing Interest Statement

The authors have declared no competing interest.

